# Endangered Przewalski’s horse, *Equus przewalskii*, cloned from historically cryopreserved cells

**DOI:** 10.1101/2023.12.20.572538

**Authors:** Ben J. Novak, Oliver A. Ryder, Marlys L. Houck, Andrea S. Putnam, Kelcey Walker, Lexie Russell, Blake Russell, Shawn Walker, Sanaz Sadeghieh Arenivas, Lauren Aston, Gregg Veneklasen, Jamie A. Ivy, Klaus-Peter Koepfli, Anna Rusnak, Jaroslav Simek, Anna Zhuk, Ryan Phelan

## Abstract

Two endangered Przewalski’s horse stallions were cloned from fibroblast cells cultured and cryopreserved in 1980. These stallions are clones of a male that lived from 1975-1998 that pedigree analyses identified as a genetically valuable male for present-day conservation breeding. This is the first time that multiple healthy clones have been produced for an endangered species.

## Main Text

We report the first clones of the endangered Przewalski’s horse, *Equus przewalskii*, or Takhi as it is called by Mongolians. The clones were produced via cross-species somatic cell nuclear transfer from donor cells cryopreserved in 1980 and using domestic horses (*Equus caballus*) as oocyte donors and surrogate mothers. The identity of the clones was verified by a microsatellite panel, and whole genome sequencing of one clone verified a match to the original donor animal (Figure 1A) with minor somatic mutations, consistent with whole genome comparisons of clones of other species^1,2^. Given the clones were produced via cross-species somatic cell nuclear transfer, it was expected that their mitochondrial haplotypes should be a mismatch to the donor, which was verified by whole mitochondrial genome analysis (Figure 1B).

**Figure 1.**
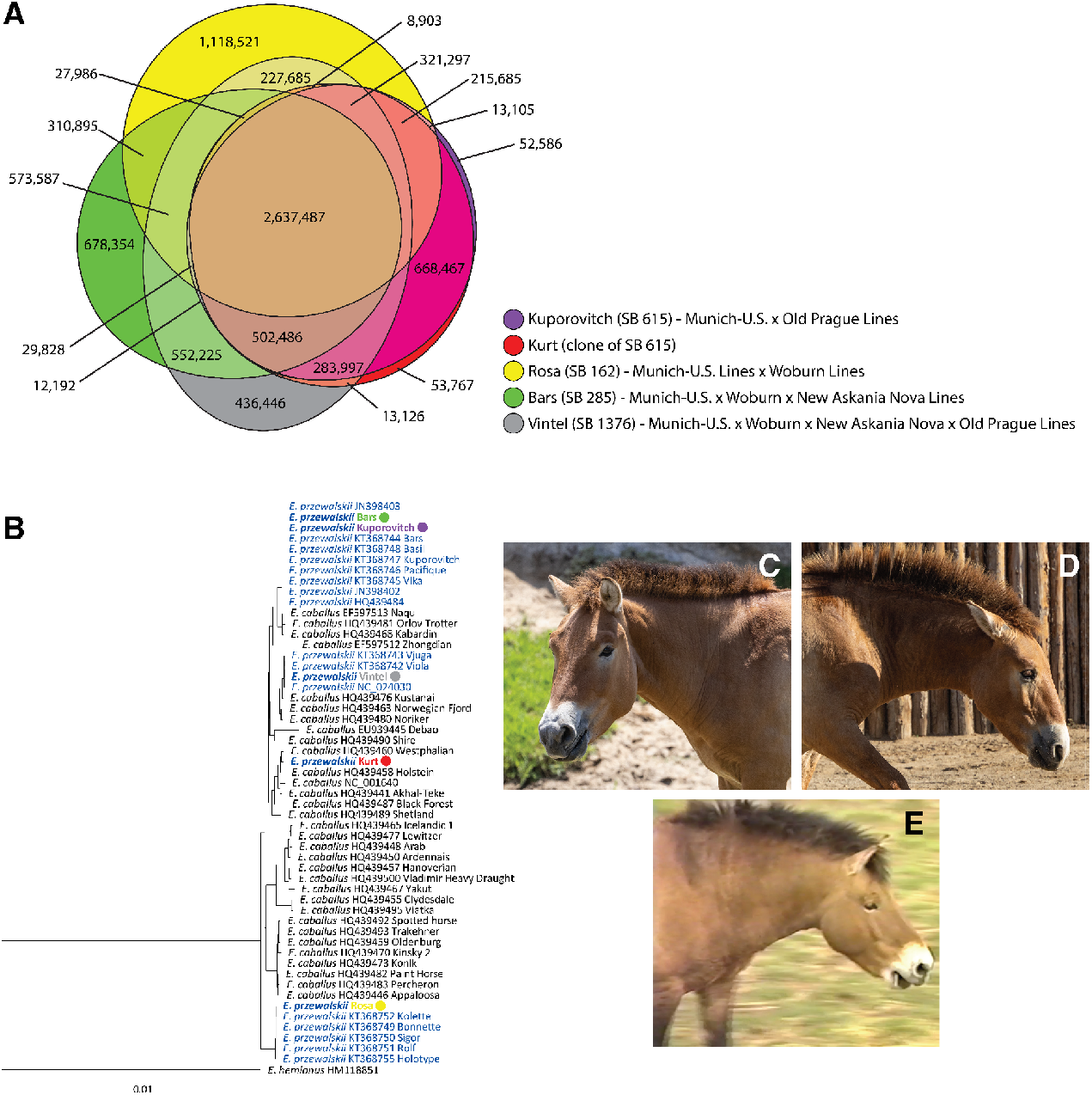
A) Euler diagram of biallelic private and shared single nucleotide variants (SNVs) for five Przewalski’s horses of multiple founder breeding lines, revealing nearly identical overlap of Kurt (clone) and Kuporovitch (donor). B) Maximum likelihood phylogenetic tree of mitochondrial genomes of domestic horses (black colored labels) and Przewalski’s horses (blue colored labels) based on an alignment of 16,130 bp. The tree is rooted with the onager (*Equus hemionus*). NCBI Genbank accession numbers are associated with domestic and Przewalski’s horses sequenced in previous studies. Asterisks indicate nodes with >90 bootstrap support values, out of 500 replicates. Przewalski’s horse individuals that were newly sequenced for this study are indicated by colored circles corresponding to A, revealing the expected mismatch between Kurt and Kuporovitch. Scale bar at bottom refers to the number of substitutions per site along branches. C) Kurt, the first clone born August 6 2020, at 25 months of age. D) Ollie, the second clone, born February 17, 2023, at 7 months of age. Photographs courtesy of San Diego Zoo Wildlife Alliance. E) Still-frame from archival footage of SB615, Kuporovitch, 8 years of age.

While several dozen domestic and wild species have been successfully cloned^3^, according to the International Union for the Conservation of Nature (IUCN) assessment and classifications of endangerment (Near Threatened, Vulnerable, Endangered, Critically Endangered, Extinct in the Wild), this is only the fourth endangered species to be successfully cloned and the first time that multiple viable clones of an endangered species have survived past the neonatal period (Table 1; note several amphibian species^4-8^ and one mammal, the cynomolgus monkey *Macaca fascicularis*^9^, that are currently assessed as endangered were not considered endangered at the time they were first cloned; and two mammals, the sand cat, *Felis margarita*^10^, and the European mouflon, *Ovis aries musimon*^11^, were erroneously considered endangered at the time of cloning but have since been reclassified).

**Table 1.** History of Cloning Endangered Species.

| Common Name | Species Name | IUCN Status | Year(s) Cloned | Number of Live Births | Number surviving neonatal period | Reproductive Capability | References |
| --- | --- | --- | --- | --- | --- | --- | --- |
| Gaur | *Bos gaurus* | Vulnerable | 2001, 2012 | 1, 1 | 0 | - | 12, 13 |
| Banteng | *Bos javanicus* | Endangered | 2003 | 2 | 1 | Infertile (cryptorchidism) | 14, 12 |
| Esfahan Mouflon | *Ovis gmelini isphahanica* | Near Threatened | 2010, 2015 | 2, NA | 0, NA | Not reported. | 15, 16 |
| Przewalski's Horse | *Equus przewalskii* | Endangered | 2020, 2023 | 2 | 2 | Not yet sexually mature. | Reported here. |

The two males produced so far, named Kurt and Ollie (Figure 1C, 1D), are clones of International Studbook Number 615 (SB615), Kuporovitch (Figure 1E), born 1975 and died 1998. Recruitment of this cell line was deemed genetically beneficial, as analyses suggested the cell line to be underrepresented within multiple regionally managed conservation breeding programs (Methods, Table 2). This is only the second species in which decades-old biobanked materials have been used to produce clones for conservation management (the first time being a Banteng, *Bos javanicus*, cloned from 25-year-old cells^14,12^, Table 1) and the first viable opportunity for clones of an endangered species to fulfill their purpose: to reproduce and rescue valuable genetic diversity for conservation breeding^17-19^. While neither of the clones has yet reached sexual maturity during a breeding season event (Przewalski’s horses reach sexual maturity at 2.5-3 years of age^20^), the testicular development of each clone is so far normal.

**Table 2:**
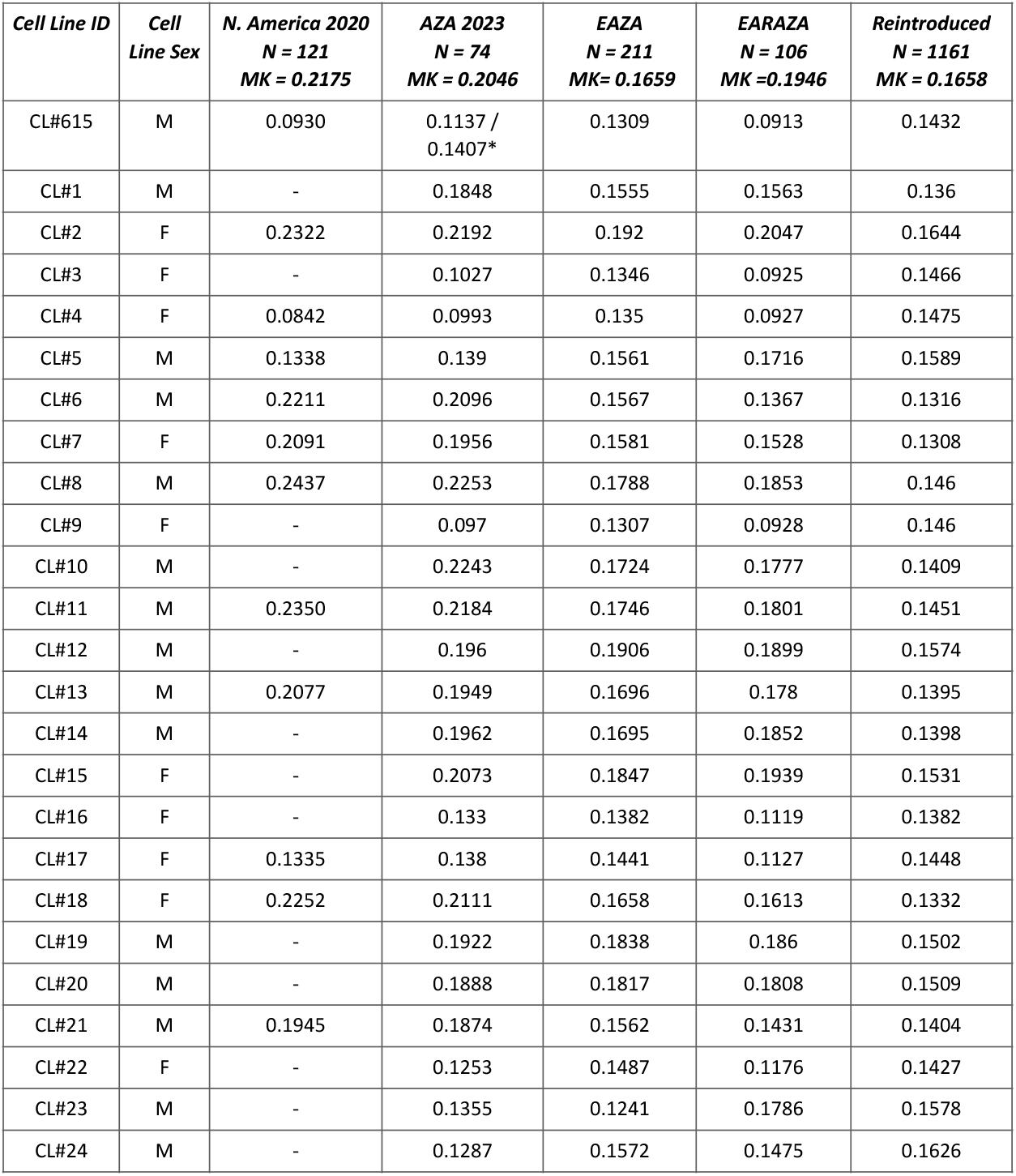
The mean kinship of each analyzed cell line (CL) relative to each population. Population size (N) and average mean kinship (MK). Cell line 615, chosen for both the 2020 and 2023 clones, is in bold. *Mean kinship of cell line 615 aver siring 7 hypothetical offspring. Identity of other cell line studbook numbers is not disclosed, as they are proprietary to multiple institutions.

| *Cell Line ID* | *Cell Line Sex* | *N. America 2020*<br>*N = 121*<br>*MK = 0.2175* | *AZA 2023*<br>*N = 74*<br>*MK = 0.2046* | *EAZA*<br>*N = 211*<br>*MK= 0.1659* | *EARAZA*<br>*N = 106*<br>*MK =0.1946* | *Reintroduced*<br>*N = 1161*<br>*MK = 0.1658* |
| --- | --- | --- | --- | --- | --- | --- |
| CL#615 | M | 0.0930 | 0.1137 / 0.1407* | 0.1309 | 0.0913 | 0.1432 |
| CL#1 | M | - | 0.1848 | 0.1555 | 0.1563 | 0.136 |
| CL#2 | F | 0.2322 | 0.2192 | 0.192 | 0.2047 | 0.1644 |
| CL#3 | F | - | 0.1027 | 0.1346 | 0.0925 | 0.1466 |
| CL#4 | F | 0.0842 | 0.0993 | 0.135 | 0.0927 | 0.1475 |
| CL#5 | M | 0.1338 | 0.139 | 0.1561 | 0.1716 | 0.1589 |
| CL#6 | M | 0.2211 | 0.2096 | 0.1567 | 0.1367 | 0.1316 |
| CL#7 | F | 0.2091 | 0.1956 | 0.1581 | 0.1528 | 0.1308 |
| CL#8 | M | 0.2437 | 0.2253 | 0.1788 | 0.1853 | 0.146 |
| CL#9 | F | - | 0.097 | 0.1307 | 0.0928 | 0.146 |
| CL#10 | M | - | 0.2243 | 0.1724 | 0.1777 | 0.1409 |
| CL#11 | M | 0.2350 | 0.2184 | 0.1746 | 0.1801 | 0.1451 |
| CL#12 | M | - | 0.196 | 0.1906 | 0.1899 | 0.1574 |
| CL#13 | M | 0.2077 | 0.1949 | 0.1696 | 0.178 | 0.1395 |
| CL#14 | M | - | 0.1962 | 0.1695 | 0.1852 | 0.1398 |
| CL#15 | F | - | 0.2073 | 0.1847 | 0.1939 | 0.1531 |
| CL#16 | F | - | 0.133 | 0.1382 | 0.1119 | 0.1382 |
| CL#17 | F | 0.1335 | 0.138 | 0.1441 | 0.1127 | 0.1448 |
| CL#18 | F | 0.2252 | 0.2111 | 0.1658 | 0.1613 | 0.1332 |
| CL#19 | M | - | 0.1922 | 0.1838 | 0.186 | 0.1502 |
| CL#20 | M | - | 0.1888 | 0.1817 | 0.1808 | 0.1509 |
| CL#21 | M | 0.1945 | 0.1874 | 0.1562 | 0.1431 | 0.1404 |
| CL#22 | F | - | 0.1253 | 0.1487 | 0.1176 | 0.1427 |
| CL#23 | M | - | 0.1355 | 0.1241 | 0.1786 | 0.1578 |
| CL#24 | M | - | 0.1287 | 0.1572 | 0.1475 | 0.1626 |

The Przewalski’s horse was extirpated from its wild range across the central Eurasian grasslands with the last commonly cited wild sighting in the Dzungarian Gobi region in 1969^21^, at which time, according to the Przewalski’s Horse International Studbook, 174 individuals survived scattered across several European and North American zoos. Those animals were descended from several founder lines that had been maintained for display. Fortunately, by the 1950s, zoo managers recognized that the future of the species relied on the horses surviving in ex situ herds. In 1959, the first International Symposium on the Preservation of the Przewalski’s Horse was organized by the Prague Zoo, initiating the first efforts to establish a plan to not only save the species from extinction but to organize a coordinated breeding program to produce enough horses to begin restoring herds to the wild. The plan was formalized at the 5th International Symposium in 1990^22^. The breeding program had been rather successful and following the 5th Symposium, reintroductions began in the early 1990s. Today, several hundred horses live in free-roaming and semi-free-roaming herds in Mongolia, China, Russia, Kazakhstan, and Hungary. The world’s ∼3,000 living Przewalski’s horses are the result of over 8,700 horses bred over nearly 20 generations from just 12 wild founders captured from 1898-1947.

While the history of the Przewalski’s horse is similar to many other species saved from extinction through conservation breeding and reintroductions, such as the California Condor, Whooping Crane, or black-footed ferret, its breeding management history is complex due to its origins preceding the modern era of zoo conservation. To illuminate by contrast, the founders of the endangered black-footed ferret breeding program were captured over a short period of time, brought into captivity together, and bred in a manner to attempt to retain as much genetic diversity as possible. The result is that all living black-footed ferrets today trace their ancestry to all founders in roughly equal proportions^23^. The Przewalski’s horse founders were captured over a long period of time, beginning in the 1890s, and scattered across several zoos in which they were bred as separate lineages for many decades before the coordinated efforts for conservation were initiated. The founders were not captured and bred with the intent to save the species nor necessarily to maximize the retention of surviving genetic diversity. Thus, when coordinated efforts for restoration began, the population was represented by five distinct lineages, two of which whose founders were interbred with domestic horses (*Equus caballus*)^24^.

Even when concerted restoration breeding efforts began, there were significant differences of opinion as to how best to interbreed the five founder lineages. A desire for “genetic purity” led to continued and problematic segregation. Combined with breeding policies aimed at purging domestic horse ancestry within introgressed lines, these management practices led to significant losses of wild Przewalski’s horse genetic diversity and high inbreeding coefficients. Experts at the time disagreed about the phenotypic impacts of the introgression of the Mongolian domestic mare, but to our knowledge, a comprehensive morphological analysis has never been reported. Fortunately, amidst this complex breeding history, initially uninformed by currently accepted practices of conservation management, biopsies of a large number of historic Przewalski’s horses were collected and cryopreserved, establishing a biobank today of nearly 600 individuals (298 individuals represented with cell lines and 277 represented by tissues preserved in DMSO from which cell lines could potentially be derived post-thaw), with several individuals just 1-5 generations separated from the wild founders.

Many individuals with domestic horse ancestry were intentionally and detrimentally underutilized in the program, including SB615, the donor sire of the clones reported here. Previous pedigree analyses concluded that significant historic allelic diversity would be lost without increasing representation of this lineage^24^, yet despite those findings SB615 was only bred during four out of 21 potential breeding seasons in his lifetime. When selecting individuals to clone for ongoing genetic management, the genetic values of available cell lines were evaluated in the context of current conservation breeding programs. The cell line from SB615 was chosen to clone because this line had the lowest mean kinship of all males, indicating a genetic benefit for recruitment (Methods, Table 2).

The reproducible success of cloning this decades-long cryopreserved cell line demonstrates the potential to clone other cryopreserved cell lines for ongoing genetic management. Thanks to the extensive biobanking efforts of cell lines and germplasm (gametes), cloning more historic individuals or “resurrecting” stallions through artificial insemination^25^ offers the chance to undo some of the genetic erosion caused by decades of managing the small zoo populations. In practice, the extensive biobank of Przewalski’s horse cell lines and gametes can now be viewed as an active reproductive resource for future breeding decisions. The use of advanced reproductive technologies is still rare in conservation, owing partially to the nascency of reproductive knowledge and techniques for most species. While biobanking of viable cells and gametes for endangered species has been established in multiple centers, its potential impacts are to this point limited by the small number of taxa cryopreserved and low degree of these collections to capture extant variation^26^, despite well-established methods for many species, particularly for mammals^27^. The Przewalski’s horse recovery program could be the first to demonstrate the ideal model system for biobanking for conservation impact^19^.

## Methods

### Pedigree Analysis

For the 2020 clone, biobanked cell lines were analyzed to determine which would confer the greatest genetic benefit to the cooperatively managed Association of Zoos and Aquarium (AZA) Przewalski’s horse population. Cooperative management for this population occurs through a strategy of minimizing pedigree-based kinships^28,29^. The mean kinship of an individual is the average kinship between that individual and all other living individuals in the population including itself. Reproduction from breeding pairs with low mean kinships and inbreeding coefficients below the population’s average mean kinship are prioritized. Twelve cell lines housed at the San Diego Zoo Wildlife Alliance Biodiversity Bank’s Frozen Zoo® were each independently analyzed to compare their mean kinships, should they be added to the population, with the average mean kinship of the managed population living at the time of analyses. The population included both AZA and non-AZA facilities that were participating in cooperative management. Data from the AZA Regional Asian Wild Horse (*Equus przewalskii*) Studbook (current to December 31, 2019^30^) was exported from Poplink 2.5.2^31^ to PMx v.1.6.20190628^32^ to calculate pedigree-based kinships. Sterilized males and females over the age of 25 were excluded from kinship analyses. Each cell line, independently of the others, was added to the genetically managed AZA population to estimate the cell line’s mean kinship within the population. The mean kinship of each cell line was then ranked from lowest mean kinship to highest mean kinship (Table 2).

The kinship analysis to prioritize cell lines for the 2023 clone were performed using the same methods as the 2020 clone, with the following modifications. Thirteen additional cell lines were added to the genetic analyses, for a total of 25 cell lines examined. Data from the International Studbook for Przewalski’s horse was exported from the ZIMS Studbook (current to December 2021^33^) to PMx v.1.7.0.20210915 to calculate pedigree-based kinships. In addition to comparing the mean kinship of the cell lines in the context of the AZA *ex situ* population that included the young 2020 clone, mean kinships of the cell lines within three other regional populations were examined: the European Association of Zoos and Aquariums (EAZA) *ex situ* population, the Eurasian Regional Association of Zoos and Aquariums (EARAZA) *ex situ* population, and the combined reintroduced free-roaming and semi-free populations of Orenburg Nature Reserve (Russia), Gobi National Park (Mongolia), Hortobágy National Park (Hungary), and Hustai Nuruu National Park (Mongolia). To account for the possible genetic representation from future reproduction of the 2020 clone within the AZA population, 8 hypothetical offspring were added to the AZA population where the 2020 clone was the sire and 7 reproductive females residing at San Diego Zoo Safari Park were the dams. The mean kinship of each cell line was then ranked from lowest mean kinship to highest mean kinship within each region (Table 2).

### Somatic Cell Nuclear Transfer

To clone SB615, a vial of the historically cryopreserved fibroblast cells was shipped from the San Diego Zoo Wildlife Alliance Frozen Zoo^®^ to the commercial cloning company ViaGen Pets & Equine’s Texas based facility where the cells were thawed and donor cells were fused with enucleated domestic horse oocytes to produce a total of 11 reconstructed embryos. The reconstructed embryos were transferred to 11 domestic horse surrogate mares (1 embryo per mare). Two ensuing pregnancies resulted in live births (Table 3, Figure 1).

**Table 3.** Cross-species cloning results for donor cell line SB615. ^*^Gestation includes 7 days *in vitro*.

| **Trial Year** | **Embryos Transferred** | **Surrogates** | **Pregnancies** | **Live Births (% total embryos)** | **Gestation Length\*** |
| --- | --- | --- | --- | --- | --- |
| 2020 | 4 | 4 | 3 (75%) | 1 (25%) | 322 days |
| 2023 | 7 | 7 | 4 (57%) | 1 (14%) | 337 days |
| Total | 11 | 11 | 7 (63%) | 2 (18%) | - |

### Microsatellite Genotyping

Blood samples were collected from the clone horses and shipped to the University of California Davis Veterinary Medicine Veterinary Genetics Laboratory for nuclear microsatellite genotyping along with a cell pellet sample of the SB615 donor cells. A panel of 15 nuclear microsatellite loci were analyzed for each clone, which include the mandatory loci for assessing equine parentage recommended by the International Society for Animal Genetics (https://www.isag.us/): *AHT4, AHT5, AME, ASB2, ASB17, ASB23, HMS2, HMS3, HMS6, HMS7, HTG10, HTG4, LEX3, LEX33, VHL20*^34^. The two clones were a match to the donor cells across all loci.

### Mitochondrial and Whole Genome Comparisons

Given the close evolutionary relationship and introgression between domestic and Przewalski’s horses, previous mtDNA studies have found shared haplotypes between the species^35,36^, and therefore a single mtDNA locus PCR may not provide adequate resolution for identity comparisons. These factors may also obscure microsatellite analyses. Therefore whole genome sequencing was performed to compare nuclear/mitochondrial DNA identities of one of the two clones to the donor cells alongside a selection of additional unrelated Przewalski’s horses as a control data set. Cell pellets from cultured fibroblasts of each horse were shipped to the commercial service lab Psomagen, Inc. (Rockville, Maryland, USA) for short-read resequencing. Specifically, genomic DNA was extracted from cell pellets using the Mag-Bind Blood and Tissue Kit (Omega Bio-Tek Inc., Norcross, GA). DNA concentrations were assessed using Picogreen and Victor x2 fluorometry (Life Technologies, Carlsbad, CA) and an Agilent 4200 Tapestation (Agilent Technologies, Santa Clara, CA), and quality was checked with 1% TBE gel electrophoresis. Genomic DNAs were then sheared into 350 bp fragments with a Covaris S220 Ultrasonicator (Covaris, Woburn, MA). DNA fragments were used to generate genomic libraries using the TruSeq DNA PCR-free library kit (Illumina, San Diego, CA), which were quality checked on an Agilent 4200 Tapestation and quantitated using a Lightcycler qPCR assay (Roche Life Science, St. Louis, MO). Libraries were paired-end sequenced (2 x 150 bp) to a depth of 20x on an Illumina NovaSeq 6000 instrument, which generated ∼56 Gb for each sample.

Paired-end reads were explored with FastQC to calculate and visualize sequence quality metrics^37^. The reads were aligned to the reference genome assembly of domestic horse (GCF_002863925.1) using the BWA-MEM-(v0.7.17) algorithm with default parameters^38^, and the reference was indexed, and a dictionary created using SAMtools (v.1.17)^39^ and Picard (v2.27.5)^40^. The mapped reads were sorted with SAMtools (v1.17), and duplicated reads were marked with Picard (v2.27.5) using MarkDuplicates. Finally, alignment statistics were evaluated with Picard and SAMtools.

SNVs were called using GATK HaplotypeCaller (v4.2.6.1)^41^, followed by the use of GATK SelectVariants to obtain only biallelic SNVs. VariantFiltration was applied to filter the vcf files based on the following parameters: “QD < 2.0” -filter-name “QD2”, “QUAL < 30.0” -filter-name “QUAL30”, “SOR > 3.0” -filter-name “SOR3”, “FS > 60.0” -filter-name “FS60”, “MQ < 40.0” -filter-name “MQ40”, -filter “MQRankSum < -12.5 “, -filter “ReadPosRankSum < -8.0”. The variant call set was further filtered by excluding multiallelic variants and setting a read depth threshold of DP < 5 using BCFtools (v1.17)^39^.

### Private Alleles Analysis

To illustrate the proportions of private SNVs and those that are shared between individuals, a Euler diagram was created based on biallelic SNVs alone. VCF files were parsed and for each individual we prepared a list of values containing SNVs and its benchmarks. Using the Eulerr R package^42^ and these lists, we then prepared a Euler diagram. The diagram showed a huge intersection across the clone (Kurt) and the donor (Kuporovitch, SB 615); from a genetic point of view the two individuals are equal. Number of alleles shared across Kurt and Kuporovitch (668,467) is comparable to numbers of private alleles in other individuals which vary from 436,446 in Vintel SB1376 to 1,118,521 in Rosa SB162 (representing the latest and earliest generations of the pedigree in our sample set, respectively). Mitochondrial genome assembly and analysis

The mitochondrial genomes of the five sequenced Przewalski’s horses were *de novo* assembled using GetOrganelle version 1.7.7.0^43^. The full read set for each individual was used as input and the complete mitochondrial genome of the domestic horse was employed as the seed, with the -F animal_mt parameter and all remaining parameters set to their default values. The mean coverage of the assemblies ranged from 342X – 1,619X, as estimated with Mosdepth^44^. We obtained mitochondrial genome assemblies ranging in length from 16,576 – 16,593 bp, with the length differences corresponding to the number of tandem repeats in the control region. The mitogenomes were annotated in the MITOS2 webserver^45^.

To assess the phylogenetic position of the five Przewalski’s horses with respect to their maternal ancestry, particularly the cloned individual Kurt, we downloaded mitochondrial genomes of available Przewalski’s horses (n = 16) as well as those from a diversity of domestic horse breeds (n = 32) from NCBI’s Genbank, which were published in previous studies^46-49^. We used the mitochondrial genome phylogeny reported by Der Sarkissian et al.^49^ to select domestic horse breeds that provided the maximal representation of mitochondrial lineage diversity. We also downloaded the mitogenome of an onager, *Equus hemionus* (HM118851^50^), which was used as the outgroup to root the phylogenetic tree. All mitogenomes were imported into Geneious Prime version 2023.2 (https://www.geneious.com), aver which a 16,748 bp multiple sequence alignment was generated using the MAFFT version 7.490 plugin with the following settings: AUTO algorithm, 200PAM / k=2 scoring matrix, 1.53 gap open penalty, and an offset value of 0.123^51^. Due to poor alignment and missing sequence data for some individuals in the control region, we trimmed 618 bp from this region, resulting in a final alignment of 16,130 bp. This alignment was used to construct a maximum likelihood phylogeny using the Geneious Prime plugin of RAxML version 8.2.11^52^. The nucleotide substitution model was set as GTRGAMMA and the search algorithm was set as “Rapid bootstrapping and search for beast-scoring ML tree”, with number of bootstrap replicates = 500, parsimony random seed = 3, and starting tree = random. The resulting tree was visualized in Geneious Prime with subsequent modifications done in Microsoft PowerPoint.

Phylogenetic analysis of mitochondrial genomes confirmed that Kurt’s maternal ancestry is derived from a domestic horse, as his mitogenome is nested within a clade of domestic horses representing Westphalian, Holstein, Akhal-Teke, Black Forest, and Shetland breeds, with >90% bootstrap support across most nodes within the clade. In contrast, the four other individuals with an expected Przewalski’s horse maternal ancestry, Bars, Kuporovitch, Rosa and Vintel, are each grouped within one of the three clades containing other Przewalski’s horse mitogenomes, which are supported by >90% bootstrap values. Notably, the new mitogenomes of Bars and Kuporovitch fall within the same haplogroup that includes the previous mitogenomes sequenced from these two individuals, KT368744 and KT368747, respectively.

## References

1. Kim, HM., Cho, Y., Kim, H. et al. Whole genome comparison of donor and cloned dogs. Sci Rep 3, 2998 (2013).

2. Jivanji, S., Harland, C., Cole, S. et al. The genomes of precision edited cloned calves show no evidence for off-target events or increased de novo mutagenesis. BMC Genomics 22, 457 (2021).

3. Klinger, B., & Schnieke, A. 25th ANNIVERSARY OF CLONING BY SOMATIC-CELL NUCLEAR TRANSFER Twenty-five years aver Dolly: how far have we come? Reproduction, 162(1), F1–F10 (2021).

4. Tung TC, Wu SC, Tung YYF, Yan SY, Tu M, Lu TY . Nuclear transplantation in fish. Science bulletin, Academia Sinica (in Chinese) 7:60–1 (1963).

5. Sambuichi, H. The roles of the nucleus and the cytoplasm in development I. An interspecific hybrid frog, developed from a combination of Rana nigro-maculata nigromaculata cytoplasm with a diploid nucleus of Rana nigromaculata brevipoda. Journal of Science, Hiroshima University Ser. B. Div. 17:33–41 (1957).

6. Signoret, J. Transplantations nucléaires et différenciation embryonnaire. Arch. Biol. 71:591–606 (1965).

7. Aimar, C. Analyse par la greffe nucléaire des propriétés morphogéné-tiques des noyaux embryonnaires chez Pleurodeles waltlii (Amphibien Urodèle). Application à l’étude de la gémellarité expérimentale. Ann. Embryol. Morphogen. 5:5–42 (1972).

8. Nishioka, M. Reciprocal nucleo-cytoplasmic hybrids between Rana brevipoda and Rana plancyi chosenica. Scientific Report of the Laboratory for Amphibian Biology, Hiroshima University. (1972).

9. Liu, Z., Y. Cai, Y. Wang, Y. Nie, C. Zhang, Y. Zu, X. Zhang, Y. Lu, Z. Wang, M. Poo, and Q. Sun. Cloning of Macaque Monkeys by Somatic Cell Nuclear Transfer. Cell 172:881–887 (2018).

10. Gómez, M. C., C. E. Pope, R. H. Kutner, D. M. Ricks, L. A. Lyons, M. Ruhe, C. Dumas, J. Lyons, M. López, B. L. Dresser, and J. Reiser. Nuclear transfer of sand cat cells into enucleated domestic cat oocytes is affected by cryopreservation of donor cells. Cloning and Stem Cells 10:469–483 (2008).

11. Loi, P., Ptak, G., Barboni, B., Fulka, J., Cappai, P., & Clinton, M. Genetic rescue of an endangered mammal by cross-species nuclear transfer using post-mortem somatic cells. Nat Biotechnol 19, 962–964 (2001).

12. Friese, C. Cloning Wildlife: Zoos, Captivity, and the Future of Endangered Animals. New York: New York University Press. (2013).

13. Srirattana, K., Imsoonthornruksa, S., Laowtammathron, C., Sangmalee, A., Tunwattana, W., Thongprapai, T., Chaimongkol, C., Ketudat-Cairns, M., & Parnpai, R.Full-term development of gaur-bovine interspecies somatic cell nuclear transfer embryos: Effect of Trichostatin A treatment. Cellular Reprogramming 14, 248–257 (2012).

14. Janssen, D. L., M. L. Edwards, J. A. Koster, R. P. Lanza, & O. A. Ryder. 206 Postnatal management of cryptorchid banteng calves cloned by nuclear transfer utilizing frozen fibroblast cultures and enculeated cow ova. Reproduction, Fertility and Development 16, 224 (2004).

15. Hajian, M., Hosseini, S. M., Forouzanfar, M., Abedi, P., Ostadhosseini, S., Hosseini, L., Moulavi, F., Gourabi, H., Shahverdi, A.H., Taghi Dizaj, A. V., Kalantari, S. A., Fotouhi, Z., Iranpour, R., Mahyar, H., Amiri-Yekta, A., & Nasr-Esfahani, M. H. “Conservation cloning” of vulnerable Esfahan mouflon (Ovis orientalis isphahanica): In vitro and in vivo studies. European Journal of Wildlife Research 57, 959–969 (2011).

16. Dehghan, S. K. Scientists in Iran clone endangered mouflon – born to domestic sheep. The Guardian. https://www.theguardian.com/science/2015/aug/05/iran-scientists-clone-endangered-mouflon-domestic-sheep. (2015).

17. Ryder, O. A. and K. Benirschke. The potential use of “cloning” in the conservation effort. Zoo Biology 16, 295–300 (1997).

18. Ryder, O.A. Cloning Advances and Challenges for Conservation. TRENDS in Biotechnology 20, 231–232 (2002).

19. Ballou, J.D., Lacy, R.C., Traylor-Holzer, K., Bauman, K., Ivy, J.A. and Asa, C. Strategies for establishing and using genome resource banks to protect genetic diversity in conservation breeding programs. Zoo Biology 42, 175–184 (2023).

20. Monfort, S. L., Arthur, N. P., & Wildt, D. E. Reproduction in the Przewalski’s Horse. In: Przewalski’s Horse: The History and Biology of an Endangered Species, ed. Boyd L., & Houpt, K.A. State University of New York Press (1994).

21. Bouman D.T., Bouman J.G. The history of Przewalski’s Horse. In: Boyd L., Houpt D.A., editors. Przewalski’s Horse: The History and Biology of an Endangered Species. State University of New York Press; Albany, NY, USA: 1994. pp. 5–38.

22. Siefert, S. Proceedings of the Fifth International Symposium on the Preservation of the Przewalski Horse. Leipzig Zoological Garden, Leipzig, Germany. (1992).

23. Marinari, P. & Lynch, C. AZA Population Analysis & Breeding and Transfer Plan: Black-footed ferret (Mustela nigripes) AZA Species Survival Plan ® Yellow Program. (2021).

24. Geyer, C.J., Thompson, E.A. and Ryder, O.A. Gene survival in the Asian wild horse (Equus przewalskii): II. Gene survival in the whole population, in subgroups, and through history. Zoo Biol., 8, 313–329 (1989). 10.1002/zoo.1430080402

25. Pukazhenthi, B.S., Johnson, A., Guthrie, H.D., Songsasen, N., Padilla, L.R., Wolfe, B.A., da Silva, M.C., Alvarenga, M.A. & Wildt, D.E. Improved sperm cryosurvival in diluents containing amides versus glycerol in the Przewalski’s horse (Equus ferus przewalskii). Cryobiology 68(2), 205–214. (2014).

26. Mooney, A., Ryder, O. A., Houck, M. L., Staerk, J., Conde, D. A., & Buckley, Y. M. Maximizing the potential for living cell banks to contribute to global conservation priorities. Zoo Biology, 1–12. (2023).

27. Houck M.L., Lear T.L. and Charter S.J. Animal Cytogenetics, Chapter 24. In: The AGT Cytogenetics Laboratory Manual, 4th ed. Arsham M., Barch M. and Lawce H., (eds), pp. 1055–1102. John Wiley & Sons, Inc., Hoboken, NJ, USA (2017).

28. Ballou JD,& Lacy RC. Identifying genetically important individuals for management of genetic diversity in pedigreed populations. In: Ballou JD, Gilpin M, Foose TJ, editors. Population management for survival and recovery. New York: Columbia Press. P. 76–111 (1995).

29. Fernandez, J., & Toro, M. A. The use of mathematical programming to control inbreeding in selection schemes. Journal of Animal Breeding and Genetics 116, 447–466 (1999).

30. Falino, A. Asian Wild Horse (Equus przewalskii) AZA Regional Studbook, Association of Zoos and Aquariums, Silver Springs, MD. (2021).

31. Faust, L.J., Bergstrom, Y.M., Thompson, S.D., & Bier, L. PopLink Version 2.5.2. Chicago, IL: Lincoln Park Zoo. (2018).

32. Lacy RC, Ballou JD, Pollak JP. (2012). PMx: Software package for demographic and genetic analysis and management of pedigreed populations. Methods in Ecology and Evolution, 3(2), 433–437.

33. ZIMS for Studbooks for Przewalski’s Horse. (Simek, J., Prague Zoo, Current to January 2022). Species360 Zoological Information Management System. Retrieved from http://zims.Species360.org

34. Bowling, A. T., Eggleston-Stom, M. L., Byrns, G., Clark, R. S., Dileanis, S., & Wictum, E. Validation of microsatellite markers for routine horse parentage testing. Animal Genetics 28(4), 247–252 (1997).

35. Goto, H., Ryder, O.A., Fisher, A.R., Schultz, B., Kosakovsky Pond, S.L., Nekrutenko, & A., Makova, K.D. A massively parallel sequencing approach uncovers ancient origins and high genetic variability of endangered Przewalski’s horses. Genome Biol Evol. 3, 1096–106 (2011).

36. Vilstrup, J.T., Seguin-Orlando, A., Stiller, M., Ginolhac, A., Raghavan, M., Nielsen, S.C.A, et al. Mitochondrial Phylogenomics of Modern and Ancient Equids. PLoS ONE 8(2), e55950 (2013).

37. Andrews, S., Krueger, F., Segonds-Pichon, A., Biggins, L., Krueger, C., & Wingett, S. FastQC. A quality control tool for high throughput sequence data, 370 (2010).

38. Li, H. and R. Durbin. Fast and accurate short read alignment with Burrows–Wheeler transform. Bioinformatics 25(14): 1754–1760 (2009).

39. Danecek, P., J. K. Bonfield, J. Liddle, J. Marshall, V. Ohan, M. O. Pollard, A. Whitwham, T. Keane, S. A. McCarthy, R. M. Davies and H. Li. Twelve years of SAMtools and BCFtools. GigaScience 10(2) giab008 (2021).

40. Broad, I. Broad Institute. Picard Tools. Broad Institute, GitHub Repository. 2021. http://broadinstitute.github.io/picard/. (2021).

41. Poplin, R., V. Ruano-Rubio, M. A. DePristo, T. J. Fennell, M. O. Carneiro, G. A. Van der Auwera, D. E. Kling, L. D. Gauthier, A. Levy-Moonshine, D. Roazen, K. Shakir, J. Thibault, S. Chandran, C. Whelan, M. Lek, S. Gabriel, M. J. Daly, B. Neale, D. G. MacArthur and E. Banks. Scaling accurate genetic variant discovery to tens of thousands of samples. bioRxiv 201178 (2018).

42. Gibson, T. Eulerr: Area-Proportional Euler and Venn Diagrams with Circles or Ellipses. R package version 6.1.1. (2021). Retrieved from https://cran.r-project.org/web/packages/eulerr/vignemes/introduction.html

43. Jin, JJ., Yu, WB., Yang, JB. et al. GetOrganelle: a fast and versatile toolkit for accurate de novo assembly of organelle genomes. Genome Biology 21, 241 (2020).

44. Pedersen, B. S., & Quinlan, A. R. Mosdepth: quick coverage calculation for genomes and exomes. Bioinformatics 34(5), 867–868 (2018).

45. Donath, A., Jühling, F., Al-Arab, M., Bernhart, S. H., Reinhardt, F., Stadler, P. F., Middendorf, M., & Bernt, M. Improved annotation of protein-coding genes boundaries in metazoan mitochondrial genomes. Nucleic Acids Research 47(20), 10543–10552 (2019).

46. Xu, S., Luosang, J., Hua, S., He, J., Ciren, A., Wang, W., Tong, X., Liang, Y., Wang, J., & Zheng, X. High altitude adaptation and phylogenetic analysis of Tibetan horse based on the mitochondrial genome. Journal of Genetics and Genomics 34(8), 720–729 (2007).

47. Lippold, S., Matzke, N. J., Reissmann, M., & Hofreiter, M. (2011). Whole mitochondrial genome sequencing of domestic horses reveals incorporation of extensive wild horse diversity during domestication. BMC Evolutionary Biology 11, 328. (2011).

48. Achilli, A., Olivieri, A., Soares, P., Lancioni, H., Hooshiar Kashani, B., Perego, U. A., Nergadze, S. G., Carossa, V., Santagostino, M., Capomaccio, S., Felicetti, M., Al-Achkar, W., Penedo, M. C., Verini-Supplizi, A., Houshmand, M., Woodward, S. R., Semino, O., Silvestrelli, M., Giulotto, E., Pereira, L., Bandelt, H-J., Torroni, A. Mitochondrial genomes from modern horses reveal the major haplogroups that underwent domestication. Proceedings of the National Academy of Sciences of the United States of America 109(7), 2449–2454 (2012).

49. Der Sarkissian, C., Ermini, L., Schubert, M., Yang, M. A., Librado, P., Fumagalli, M., Jónsson, H., Bar-Gal, G. K., Albrechtsen, A., Vieira, F. G., Petersen, B., Ginolhac, A., Seguin-Orlando, A., Magnussen, K., Fages, A., Gamba, C., Lorente-Galdos, B., Polani, S., Steiner, C., Neuditschko, M., Jagannathan, V., Feh, C., Greenblatt, C. L., Ludwig, A., Abramson, N. I., Zimmermann, W., Schafberg, R., Tikhonov, A., Sicheritz-Ponten, T., Willerslev, E., Marques-Bonet, T., Ryder, O. A., McCue, M., Rieder, S., Leeb, T., Slatkin, M., Orlando, L. Evolutionary Genomics and Conservation of the Endangered Przewalski’s Horse. Current Biology 25(19), 2577–2583 (2015).

50. Luo, Y., Chen, Y., Liu, F., Jiang, C., & Gao, Y. Mitochondrial genome sequence of the Tibetan wild ass (Equus kiang). Mitochondrial DNA, 22(1-2), 6–8 (2011).

51. Katoh, K., & Standley, D. M. MAFFT Multiple Sequence Alignment Software Version 7: Improvements in Performance and Usability. Molecular Biology and Evolution 30(4), 772–780 (2013).

52. Stamatakis, A. RAxML version 8: A tool for phylogenetic analysis and post-analysis of large phylogenies. Bioinformatics 30(9), 1312–1313 (2014).

